# Computational modeling of pro-inflammatory cytokine-enhanced blood coagulation

**DOI:** 10.64898/2026.05.02.722421

**Authors:** Geli Li, Galit H. Frydman, He Li

## Abstract

The interplay between inflammation and coagulation is a central driver of thrombotic risk across various diseases. While mathematical models of blood coagulation are well established, there remains a critical gap in quantitative frameworks that capture inflammation-induced hypercoagulability. In this study, we develop a mathematical model that explicitly simulates the interaction between pro-inflammatory cytokines and the coagulation cascade. The model incorporates key mechanisms, including: (i) upregulation of tissue factor (TF) by IL-1*β*, IL-6, and TNF-*α*; (ii) suppression of natural anticoagulants, namely antithrombin III (ATIII) and tissue factor pathway inhibitor (TFPI), by IL-6 and TNF-*α*; and (iii) feedback amplification of proinflammatory cytokines by thrombin. By encoding the bidirectional feedback between inflammatory and coagulation pathways, the model captures essential features of inflammation-driven hypercoagulability and enables systematic quantification of how variability in inflammatory extent and duration results in heterogeneous thrombin generation (TG) dynamics. To evaluate its effectiveness, we integrate the model with TG assays and apply it to virtual patient cohorts representing four clinically distinct conditions: COVID-19, sickle cell disease (SCD), type 2 diabetes mellitus (T2DM) and Hemophilia A. Model simulations predict that disease-specific inflammatory environments induce distinct shifts in TG dynamics. In COVID-19 and T2DM, elevated cytokine levels lead to shortened lag times and increased thrombin peak, whereas in SCD, shortened lag times are accompanied by a reduced thrombin peak. These effects are strongly modulated by both cytokine concentration and duration of exposure. These results demonstrate that the proposed computational model augments conventional TG assays by mechanistically linking inflammatory signaling to disease-specific coagulation responses. Collectively, the proposed computational framework extends conventional TG assays by considering the interplay between inflammation and coagulation, thereby providing a potential tool for predicting disease progression and identifying disease-specific therapeutic targets to advance personalized management strategies in thrombo-inflammatory disorders.

## 1 Introduction

Human blood coagulation is a highly orchestrated biochemical cascade that protects the body from excessive bleeding following vascular injury [1]. Its central purpose is to limit blood loss by rapidly forming a clot at the site of damage. Hemostasis, the physiological process that governs this response, relies on a coordinated interplay among three major components: vascular regulation, activation of the coagulation cascade, and fibrinolytic dissolution of the clot. Proper balance among these mechanisms is essential for maintaining homeostasis. When this balance is disrupted, serious and potentially life-threatening conditions can arise. Insufficient levels or activity of coagulation factors, as seen in hemophilia A, or dysregulated fibrinolysis [2], can result in uncontrolled bleeding [3]. Conversely, excessive production of procoagulant or prothrombotic factors can drive inappropriate clot formation, leading to thrombosis [4]. Because both bleeding disorders and thrombotic events pose significant health risks, tight regulation of hemostasis is essential for maintaining physiological stability.

Inflammation is increasingly recognized as one of the key drivers of prothrombotic events across a wide range of pathological conditions [5, 6]. Inflammatory signaling perturbs hemostatic homeostasis by simultaneously upregulating procoagulant pathways and weakening endogenous anticoagulant and fibrinolytic safeguards [7, 8]. Mechanistically, pro-inflammatory cytokines and innate immune activation promote tissue factor (TF) expression in monocytes and endothelial cells, enhance platelet activation, and amplify thrombin generation (TG), thereby accelerating the initiation and propagation of clots [9–11]. In parallel, inflammation suppresses major anticoagulant pathways, including antithrombin III (ATIII), the protein C system, and the tissue factor pathway inhibitor (TFPI), and attenuates fibrinolysis (e.g., through impaired plasmin generation and increased fibrinolytic inhibition), further shifting the system toward a hypercoagulable state [12]. When excessive or sustained, this immunothrombosis could become maladaptive, driving microvascular thrombosis, end-organ hypoperfusion, and organ dysfunction in settings, such as sepsis, severe viral infection, and chronic inflammatory disease [13, 14].

Key inflammatory mediators, including tumor necrosis factor-*α* (TNF-*α*), interleukin-6 (IL-6), and interleukin-1*β* (IL-1*β*), can directly remodel the hemostatic phenotype of vascular and immune compartments. These cytokines induce TF expression in monocytes and endothelial cells and promote a procoagulant endothelial switch, providing a potent trigger for TF-dependent TG [15, 16]. Moreover, inflammation enhances endothelial dysfunction and platelet activation, and strengthens platelet–leukocyte/platelet–innate immune cell interactions, thereby amplifying thrombin formation through coupled cellular and plasma-protease mechanisms [17–20]. Such inflammation-driven coagulation abnormalities, often referred to as thromboinflammation/immunothrombosis, are widely observed in severe inflammatory diseases, including sepsis and COVID-19, where pathways linked to TF, platelets, and neutrophils contribute to systemic coagulopathy and thrombotic complications [21–23]. However, the dynamic, nonlinear, and feedback-rich nature of inflammation–coagulation crosstalk is not adequately captured by isolated or static biomarkers (e.g., single time-point cytokines, D-dimer, PT/aPTT). This limitation motivates quantitative and systems-level approaches, including mechanistic/QSP modeling of coagulation networks, to integrate multi-factor interactions and characterize inflammation-induced prothrombotic states in a time-resolved manner [24].

Current mathematical models for simulating blood coagulation are categorized into two main types: ordinary differential equation (ODE) models and partial differential equation (PDE) models. The ODE models, designed to mimic thrombin and fibrin generation assays, operate on the principle that biochemical reactions can be represented as kinetic equations derived from experimental data [25–32]. These models focus on temporal changes, illustrating how concentrations of coagulation factors evolve over time under spatially uniform conditions. They are particularly useful for predicting concentration changes and identifying new reaction mechanisms, especially when there is a discrepancy between model predictions and actual data. In contrast, PDE-based models and other hybrid models are utilized to simulate thrombus growth, taking into account both unstirred systems and blood flow conditions [33–44]. These spatio-temporal models are essential for understanding clot growth within blood vessels, as they consider spatial concentration variations and can integrate blood flow velocity, often through a convection term, to provide deeper insights into the clot formation and growth dynamics [43, 44].

Despite significant advances in modeling coagulation pathways, existing mathematical frameworks remain insufficient to capture the complexity of inflammation-driven hypercoagulability. To address this gap, we propose a computational model that mechanistically links pro-inflammatory cytokines to coagulation factor dysregulation. Specifically, the model incorporates three clinically relevant interactions: 1) upregulation of TF expression by IL-1*β*, IL-6, and TNF-*α*; 2) cytokine-mediated suppression of the natural anticoagulants ATIII and TFPI by TNF-*α* and IL-6; and 3) thrombin-driven amplification of TNF-*α* and IL-6, reinforcing the thromboinflammatory feedback loop. We demonstrate the effectiveness of the proposed model by applying it to enhance TG assays across virtual patient cohorts representing four distinct disease states: COVID-19 coagulopathy, sickle cell disease (SCD), type 2 diabetes mellitus (T2DM), and hemophilia A. By rigorously quantifying how the pro-inflammatory cytokines alter coagulation dynamics across different hypercoagulable states, this work establishes a mechanistic framework for dissecting thromboinflammatory pathophysiology, with potential applications for predicting progression of inflammation-induced hypercoagulability and identifying disease-specific therapeutic targets to advance precision-medicine approaches to the management of thromboinflammatory disorders.

## 2 Data and Methods

### 2.1 Model structure

The model is organized into two functionally distinct but tightly coupled subsystems representing inflammation and coagulation. As illustrated in Fig. 1, the inflammation subsystem governs the time-dependent variation of IL-6, IL1-*β*, and TNF-*α*, while the coagulation subsystem captures TF-initiated TG. Within the model, inflammatory activity acts upstream of coagulation by promoting TF availability and attenuating endogenous anticoagulant regulation. Specifically, IL-6, IL1-*β*, and TNF-*α* activated monocytes and endothelial cells, thereby enhancing TF expression that triggers initiation of the coagulation cascade. On the other hand, pro-inflammatory cytokines also could suppress the natural anticoagulants, including ATIII and TFPI, shifting the hemostatic balance toward a prothrombotic state. The coagulation subsystem generates thrombin as a central output, which could subsequently activate platelets and promote the generation of a cross-linked fibrin network, two key components of thrombosis. In addition, thrombin feeds back into the inflammatory subsystem by promoting the production of IL-6, IL1-*β*, and TNF-*α*. This feedback establishes a positive amplification loop in which inflammation enhances TG while thrombin, in turn, reinforces inflammatory signaling. Such bidirectional coupling provides a mechanistic framework for simulating sustained thromboinflammatory activation.

**Figure 1:**
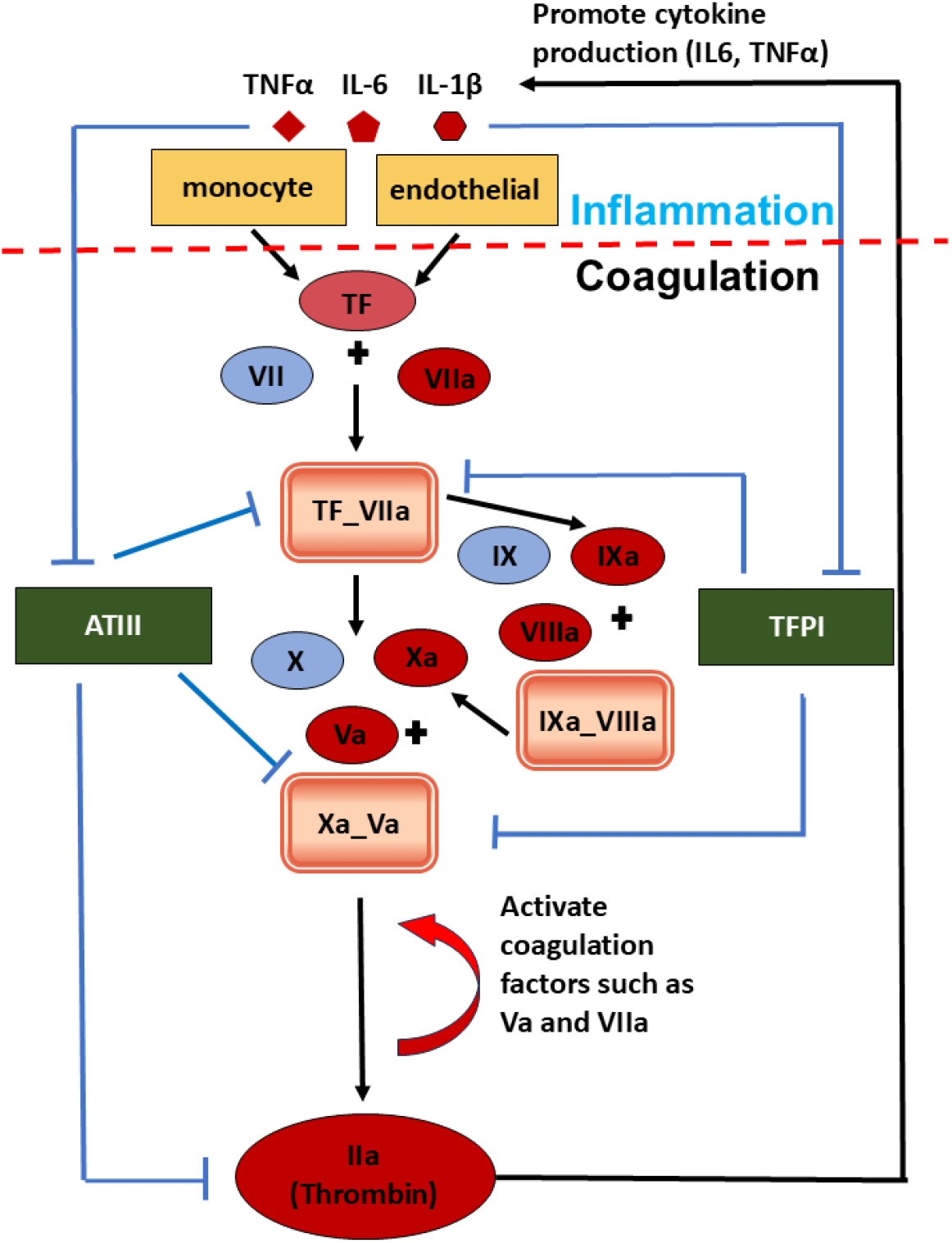
Schematic diagram of the model structure for simulating the interaction between inflammation cytokines and blood coagulation.

The coagulation module is built upon the mechanistic framework of Hockin et al. [29], which describes the core TF-initiated coagulation cascade and remains one of the most widely validated ODE-based models of hemostasis. The inflammation module incorporates three key proinflammatory cytokines, namely, IL-6, IL-1*β*, and TNF-*α*, whose interactions with coagulation factors form the mechanistic basis of thromboinflammatory coupling. The molecular pathways linking inflammatory mediators to TF upregulation, primarily through monocyte and endothelial cell activation [45, 46], and those linking thrombin to cytokine amplification via PAR-1/NF-*κ*B signaling [47, 48] are inherently intracellular, involving complex transcriptional and post-translational regulation that cannot be resolved at the level of a systems-scale ODE model. We therefore adopt a phenomenological modeling strategy, constructing quantitative input-output relationships between: 1) cytokine concentrations (IL-1*β*, IL-6, TNF-*α*) and TF production, and 2) thrombin concentration and TNF-*α*/IL-6 amplification. These relationships are parameterized directly from published experimental data, ensuring biological feasibility without requiring explicit representation of intra-cellular signaling intermediates.

To determine the production and degradation rates of each inflammatory species, we leveraged experimentally reported half-life values: IL-1*β* and IL-6 each carry a plasma half-life of approximately 1 hour [49, 50], while TNF-*α* is cleared considerably faster, with a half-life of only 4.6 minutes [51]. First-order metabolic degradation rates were derived directly from these half-lives. Production rates were then computed by imposing a steady-state constraint: in the absence of disease or external perturbation, each species must be maintained at its reported normal plasma concentration. This homeostatic calibration ensures that the model is physiologically grounded at baseline and that any simulated perturbations reflect disease-specific deviations from normal hemostatic and inflammatory tone, rather than artifacts of parameter initialization.

Overall, the proposed mathematical model simulates 38 reactions involving a total of 44 species and 72 parameters. Details of the model reaction and parameters are included in the supplementary materials. All model data, including reaction rules, species, reaction rates, and parameter values, were processed using MATLAB’s Simbiology Toolbox (MathWorks, Natick, MA). Since our model simulates a variety of cytokine and coagulation stimulation conditions, we converted all dosing units used in *in vitro* experiments (often in ng/ml) into mol per liter. ImageJ software (NIH) [52] and Getdata were utilized in the analysis and quantitative digitization of Western blot and other experimental datasets.

### 2.2 Clinical data for creating virtual patients

We generated virtual patient cohorts to investigate how the extent and duration of inflammation alter TG dynamics across four distinct diseases: COVID-19–associated coagulopathy, SCD, T2DM, and hemophilia A. Clinical laboratory ranges for coagulation factors and inflammatory markers were collected for each disease group, as summarized in Table 1. To construct these cohorts, we employed Latin hypercube sampling to efficiently explore parameter spaces and generate physiologically diverse virtual patients. Using disease-specific distributions (mean and standard deviation) of coagulation factors and cytokines as reference values, we generated 2,000 virtual patients per condition. We further calibrated the sampled parameter sets using key TG metrics, namely lag time, time to peak, and peak thrombin, derived from experimental TG data for each disease group [53–56]. This calibration step ensured that the simulated patients exhibited physiologically consistent coagulation profiles aligned with observed TG dynamics. Finally, using the TG curve as a quantitative biomarker, we systematically analyzed how pro-inflammatory cytokines modulate TG dynamics across the four disease states, thereby elucidating disease-specific thrombo-inflammatory mechanisms.

**Table 1:** Range of coagulation and cytokine factors for Normal subjects and subjects under four diseased conditions.

|  | **COVID-19** | **T2DM** | **SCD** |
| --- | --- | --- | --- |
| IL-6 | 232(1 – 32768)pg/ml [57] | 4.1 (2.3 – 5.6) pg/mL [58] | 60± 7pg/ml [59] |
| TNFα | 25(1 – 1000)pg/ml [57] | 3.8 (3.0 – 5.0) pg/mL [58] | 122.3± 16.3pg/ml [60] |
| IL1 $\beta$ | 0.8(0 – 32)pg/ml [57] | 0.2 (0.2 – 0.3) pg/mL [58] | 3.5 (0.0 – 27.26) pg/mL [61] |
| VII | 0.5mg/L (55%–170%) [62] | 0.5mg/L (60% – 130%) [63] | 0.31mg/L(±7.5%) [64] |
| VIIa | 0.2mg/L (60% – 130%) [63] | 0.2165mg/L±26.72% [65] | 0.2mg/L (60% – 130%) [63] |
| IX | 7.5mg/L ±50.6% [62] | 5mg/L (60% – 130%) [63] | 5mg/L (60% – 130%) [63] |
| X | 11.1mg/L ±27.9% [62] | 10mg/L (60% – 130%) [63] | 6.6mg/L ±10% [64] |
| II | 107.5 mg/L ±31.2% [62] | 100mg/L (60% – 130%) [63] | 75 mg/L ±10% [64] |
| VIII | 0.262 mg/L±102.9% [62] | 0.1mg/L (50% – 150%) [63] | 0.1mg/L (50% – 150%) [63] |
| V | 11.7mg/L ±44.9% [62] | 10mg/L (60% – 130%) [63] | 7.25mg/L ±17.5% [64] |
| TFPI | 70ng/mL [66] | 197.56 ± 94.88(pg/ml) [65] | 70ng/mL [66] |
| ATIII | 0.134mg/L ±15.1% [62] | 0.15mg/L [67] | 0.15mg/L [67] |
|  | **Hemophilia** | **Normal** |  |
| IL-6 | - | - |  |
| TNFα | - | - | |
| IL1 $\beta$ | - | - | |
| VII | 0.5mg/L (60% – 130%) [63] | 0.5mg/L (60% – 130%) [63] |  |
| VIIa | 0.2mg/L (60% – 130%) [63] | 0.2mg/L (60% – 130%) [63] |  |
| IX | 5mg/L (60% – 130%) [63] | 5mg/L (60% – 130%) [63] |  |
| X | 10mg/L (60% – 130%) [63] | 10mg/L (60% – 130%) [63] |  |
| II | 100mg/L (60% – 130%) [63] | 100mg/L (60% – 130%) [63] |  |
| VIII | < 1% [68] | 0.1mg/L (50% – 150%) [63] |  |
| V | 10mg/L (60% – 130%) [63] | 10mg/L (60% – 130%) [63] |  |
| TFPI | 70ng/mL [66] | 70ng/mL [66] |  |
| ATIII | 0.15mg/L [67] | 0.15mg/L [67] |  |

## 3 Results

### 3.1 Model calibration

First, we calibrate the coagulation module independently against both reference model simulations and experimental TG data under multiple perturbation conditions as reported in [29]. Figs. 2(A–C) compare thrombin dynamics simulated by the coagulation module (red curves) with those generated by a reference coagulation model under the same 25pM TF stimulations (blue circles). Specifically, thrombin responses were evaluated in the presence of ATIII and TFPI (Fig. 2(A)), in the absence of ATIII (Fig. 2(B)), in the absence of TFPI (Fig. 2(C)), in the absence of both ATIII and TFPI (Fig. 2(D)), respectively. Across all conditions, the coagulation module closely reproduced the results of the reference model, capturing key dynamical features including thrombin initiation timing, peak amplitude, and sustained or decaying response profiles. In particular, perturbations to anticoagulant pathways produced consistent qualitative and quantitative effects, indicating that the coagulation module preserves the essential regulatory structure of the reference framework [29].

**Figure 2:**
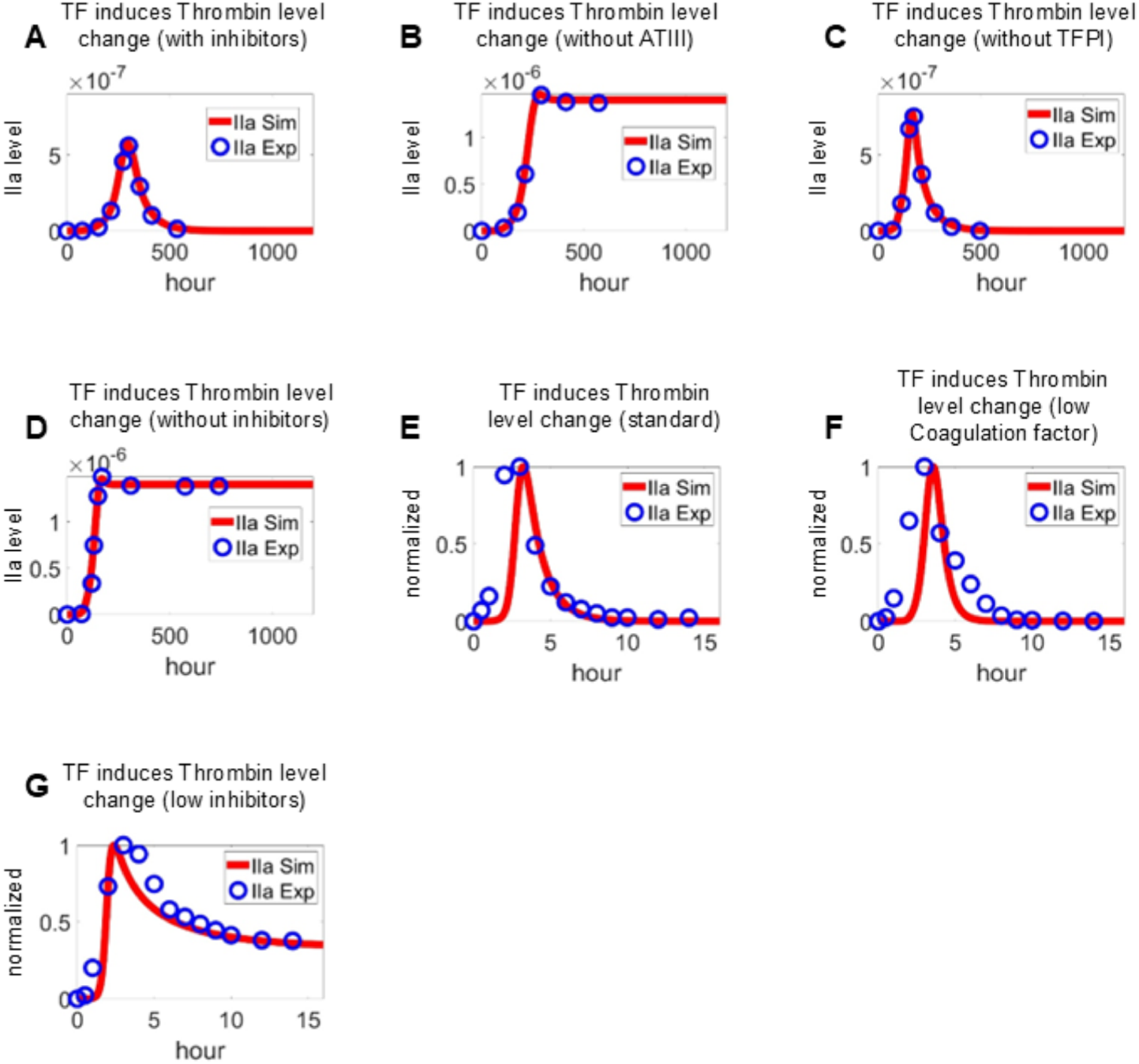
Calibration of coagulation module using TF–driven TG curves under various inhibitory conditions. TG induced by TF in the presence of ATIII and TFPI (A), in the absence of ATIII (B), in the absence of TFPI (C), in the absence of both ATIII and TFPI (D). Experimental data used in (A-D) is adopted from [29]. TG initiated by factor VIIa-TF in the presence of all proteins at mean plasma concentrations (E), in the presence of procoagulants (prothrombin, factors V, VIII, IX, and X) at 150% and anticoagulants (AT-III and TFPI) at 50% of their mean plasma concentrations (F), and in the presence of procoagulants at 50% and anticoagulants at 150% of their mean plasma concentrations (G). Experimental data used in (E-G) is adopted from [69].

To evaluate the robustness and physiological fidelity of the model, we also simulated TG dynamics under conditions corresponding to separate experimental measurements in normal plasma reported by [69]. Model predictions were compared against three experimental scenarios reflecting systematic perturbations in procoagulant and anticoagulant factor levels. Thrombin generation was initiated with 5 pmol/L factor VIIa–TF in the presence of all coagulation proteins under the following conditions: (1) baseline, with all factors at mean plasma concentrations; (2) a procoagulant-dominant state, with elevated levels (150%) of prothrombin and factors V, VIII, IX, and X, combined with reduced levels (50%) of natural anticoagulants, including ATIII and TFPI; and (3) an anticoagulant-dominant state, with reduced procoagulant levels (50%) and elevated anticoagulant levels (150%). As shown in Fig. 2(E), the model accurately reproduced the characteristic thrombin burst observed under physiological conditions, followed by a rapid decay phase. In the anticoagulant-dominant setting, both simulations and experimental data demonstrated delayed onset and attenuated peak thrombin generation (Fig. 2(F)), consistent with impaired propagation of the coagulation cascade. Conversely, in the procoagulant-dominant condition, reduced inhibitory capacity led to sustained thrombin activity with prolonged decay kinetics (Fig. 2(G)). Overall, the close agreement between simulated and experimental TG profiles across these perturbations supports the model’s ability to capture the dynamic balance between procoagulant drivers and endogenous inhibitors.

Next, we calibrated the interplay between inflammation and coagulation modules using extensive experimental datasets to constrain parameters governing the interactions between cytokines and the coagulation factors. Specifically, cell-type–specific data were used to calibrate the how the IL-6 (Figs. 3(A and B)) [70, 71]), IL-1*β* (Figs. 3(D and E)) [72, 73] and TNF-*α* (Figs. 3(F and G)) [74, 75] enhance the expression of TF on monocytes and endothelium. Furthermore, motivated by experimental observations that excessive inflammatory signaling can suppress endogenous anticoagulant pathways [76, 77], we incorporated cytokine-mediated inhibition of ATIII and TFPI into the calibration process, as shown in Figs. 3(C and H). This negative regulatory mechanism was included to capture the procoagulant shift observed under high inflammatory burden. Moreover, the activating effect of thrombin on cytokine production was calibrated using experimental data obtained at different thrombin concentrations as shown in Figs. 3 (I-J) [78], enabling the model to represent feedback amplification from coagulation and inflammatory signaling.

**Figure 3:**
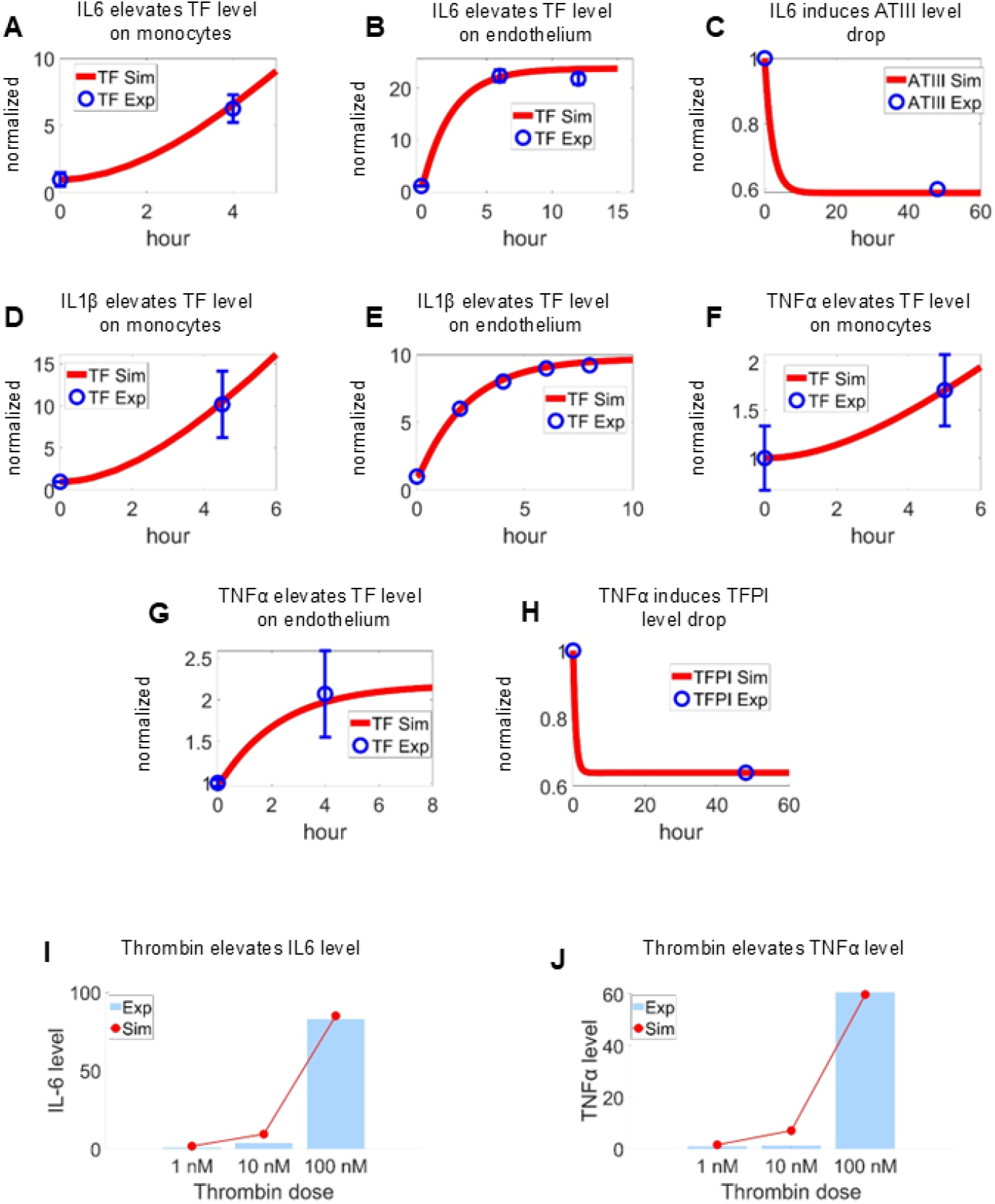
Calibration of inflammation–coagulation coupling mechanisms. (A–C) IL-6–induced TF upregulation and ATIII suppression over time [70, 71, 76]. (D–E) IL-1*β*–induced TF expression dynamics [72, 73]. (F–H) TNF-*α*–induced TF upregulation and TFPI suppression [74, 75, 77]. (I–J) Thrombin-induced cytokine production, showing dose-dependent induction of IL-6 and TNF-*α* [78]. Model simulations (red lines) were calibrated against experimental measurements (blue markers), with all outputs normalized to baseline levels.

### 3.2 Quantify the impact of pro-inflammatory cytokines on coagulation

To investigate the effects of varying inflammation intensities on coagulation, we employed two complementary biomarkers: the thrombin–antithrombin complex (ATIII-IIa) and the TG curve. The TG curve was used to characterize the kinetics of TG under cytokine-mediated perturbation, while generation of ATIII-IIa reflects the net amount of thrombin generation. Inflammatory stimulation was modeled by varying the concentrations of key pro-inflammatory cytokines at 0.5, 1, 2, 5, 10, and 50 times of their baseline values, spanning the range of cytokine levels documented in the clinical literature (Table 1). These variations represent a physiologically and clinically relevant spectrum of inflammatory states, from sub-physiologic to severely hyperinflammatory conditions. Notably, since the majority of COVID-19 cohort studies report a maximum IL-1*β* elevation of approximately 10-fold, IL-1*β* concentrations were capped at a 10× increase in all simulations to maintain clinical relevance.

Because coagulation and inflammation operate on distinct timescales: coagulation cascades unfold within seconds to minutes, whereas cytokine-mediated modulation of coagulation factor expression and endothelial activation evolves over hours, we modeled inflammatory impact on coagulation factors over 12-hour and 24-hour periods prior to initiating the coagulation module. This approach allowed sufficient time for inflammatory mediators to activate monocytes and endothelial cells, upregulating TF expression and thereby modulating the extent of thrombin generation.

In the simulations, we initiated thrombin generation with 5 pmol/L TF, consistent with established experimental conditions, to define a baseline coagulation response. Under this setting, TG was computed in the absence of pro-inflammatory cytokine modulation, providing a baseline against which the effects of proinflammatory cytokines on coagulation dynamics could be systematically evaluated. Fig. 4(A) displays the time-course of ATIII-IIa complex accumulation across cytokine doses (1×–50×), where all conditions follow a sigmoidal accumulation pattern, with a sharp transition occurring between 750–1,000 seconds. Higher cytokine concentrations produce a modest but consistent upward shift in plateau ATIII-IIa levels, suggesting that inflammation augments total thrombin generation without dramatically altering the kinetic signature. It is noted that the 1× and 2× curves are nearly superimposable, implying that a minor to mild inflammation does not meaningfully perturb net thrombin production. Quantification of the peak ATIII-IIa as a function of cytokine dose in Fig. 4(B) shows that the peak ATIII-IIa rises non-linearly, with the most dramatic increase occurring between 10× and 50×, suggesting a procoagulant tipping point at severe hyperinflammation. The relatively flat response between 1×–10× implies that mild to moderate inflammatory states, such as at the early stage of COVID-19 or sepsis, may not substantially elevate ATIII-IIa alone, potentially limiting its sensitivity as an early biomarker at lower inflammation intensities.

**Figure 4:**
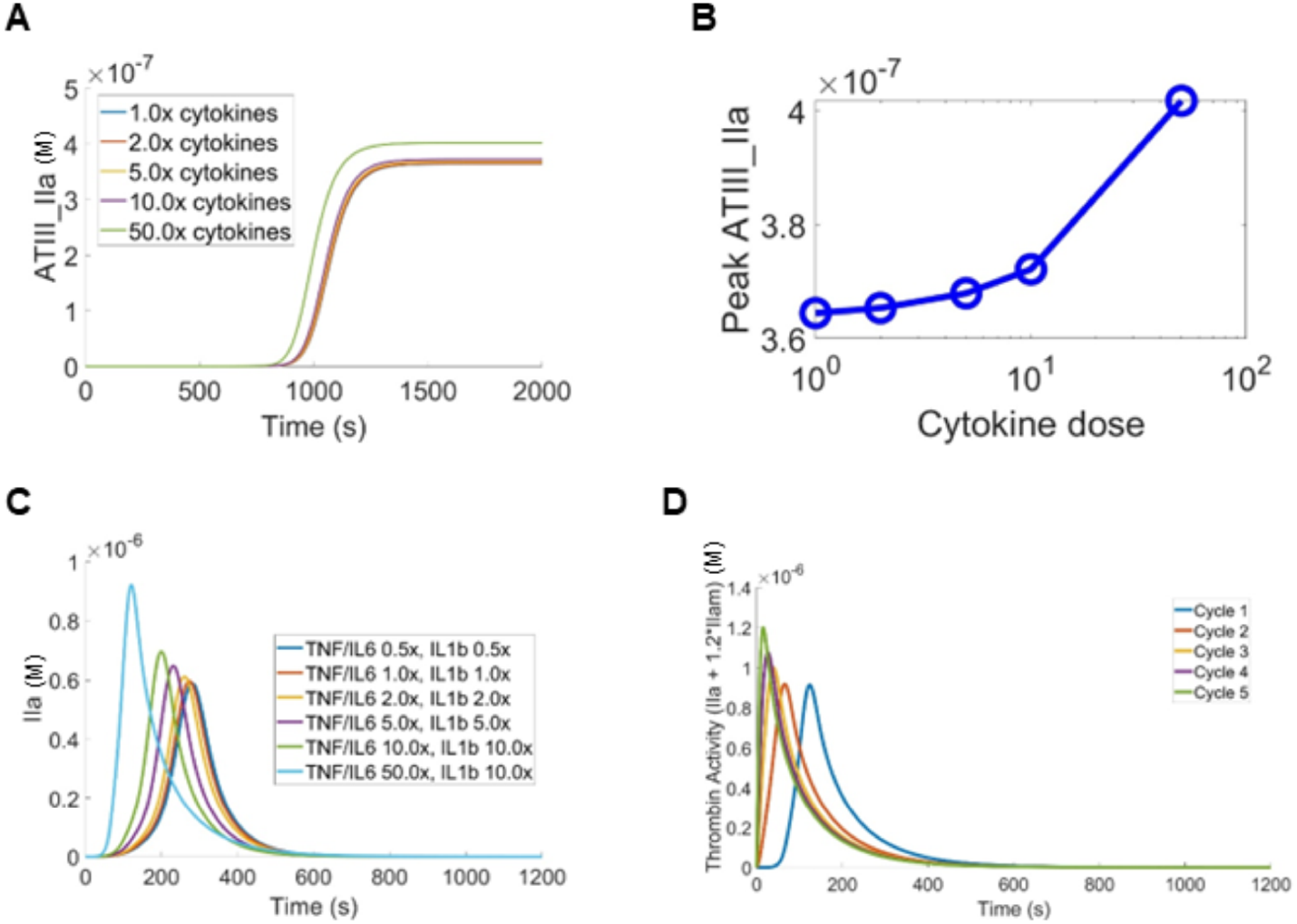
Simulated effects of pro-inflammatory cytokines on coagulation dynamics. (A) The dynamics of ATIII-IIa generation by different levels of cytokines. (B) The total amount of ATIII-IIa generated by different levels of cytokines. (C) The TG dynamics at different levels of cytokines. In (A-C), cytokine concentrations were scaled relative to the normal baseline condition. (D) Sustained inflammatory exposure at the baseline condition further enhanced thrombin activity, indicating time-dependent reinforcement of inflammation–coagulation coupling.

Fig. 4(C) illustrates the temporal dynamics of thrombin generation across the full range of cytokine doses, revealing a clear dose-dependent response. As cytokine concentrations increase from 0.5× to 10×, peak thrombin levels rise progressively. At higher inflammatory stimulation (TNF-*α* and IL-6) at 50×, thrombin increases sharply, highlighting the amplifying effect of severe inflammation on coagulation potential. Meanwhile, the time-to-peak shortens with increasing cytokine dose, indicating accelerated initiation and propagation of coagulation. This behavior is mechanistically consistent with cytokine-driven upregulation of TF expression on activated monocytes and endothelial cells, a key pathway linking inflammation to enhanced thrombin generation.

### 3.3 Simulate the amplifying cycles of inflammation and coagulation

Thrombin generation from coagulation exerts a positive feedback effect on inflammatory mediator production via PAR-1/NF-*κ*B signaling [47, 48], perpetuating the inflammation–thrombosis cycle. To capture this cyclical behavior, simulations were structured as iterative two-phase sequences: the inflammation module was executed for either 12 hours, representing sustained inflammatory states respectively, followed by a 1,200-second coagulation module. Upon completion of each coagulation phase, the inflammation module was reinitiated using the thrombin exposure metric derived from that cycle, allowing inflammatory mediator concentrations to evolve in response to the coagulation output. Coagulation factor concentrations were restored to their initial values between cycles to establish a defined computational baseline. This two-phase framework enabled systematic analysis of how inflammation-driven hypercoagulation manifests across patient cohorts with differing extent and duration of inflammation.

Fig. 4(D) shows thrombin activity across five continuous inflammation–coagulation cycles under sustained inflammatory conditions, and the results show that thrombin activity is progressively reinforced over successive cycles, indicating a cumulative amplification effect driven by prolonged inflammatory stimulation. This pattern suggests that sustained or unresolved inflammation, as opposed to a single acute inflammatory insult, is the critical determinant of escalating coagulation activity, consistent with the clinical trajectory observed in severe COVID-19 and sepsis, where persistent cytokine elevation correlates with disease worsening [9, 79, 80]. The multi-cycle amplification supports the concept that each inflammation-coagulation cycle drives the system to a more procoagulant response in the subsequent cycle, through upregulation of TF on monocytes and endothelial cells, sustained elaboration of pro-inflammatory cytokines, and suppression of endogenous anticoagulant pathways. Clinically, these results could mimic the self-reinforcing thromboinflammatory loop that underlies microvascular thrombosis and organ dysfunction in hyperinflammatory states, underscoring why early interruption of the inflammatory stimulus, rather than anticoagulation alone, may be essential to breaking the cycle [19, 81].

### 3.4 Enhance TG assays across various virtual patient cohorts

In this section, we employ the proposed model to enhance TG assays across virtual patient cohorts representing four distinct disease states: COVID-19 coagulopathy, SCD, T2DM, and hemophilia A. We first apply the coagulation factor ranges listed in Table 1 to the coagulation module and compare four commonly used metrics derived from the generated TG curves, namely lag time, time-to-peak (TTP), peak thrombin concentration, and endogenous thrombin potential (ETP) against experimentally reported values for COVID-19 coagulopathy [53], SCD [55], T2DM [54], and hemophilia A [56]. We observe that the model predictions do not agree with the TG dynamics for any of the four examined disease conditions. These findings echoed with previous studies assessing the performance of existing *in silico* coagulation models for predicting TG in normal subjects, which indicate that calibration of the coagulation module is necessary before it can reliably reproduce the key TG metrics.

To calibrate the model effectively and efficiently, we first perform a sensitivity analysis using the Latin Hypercube Sampling (LHS) method to compute Partial Rank Correlation Coefficient (PRCC) values across the coagulation factor variation ranges from the four disease groups as listed in Table 1, following the algorithm described by Marino et al. [82]. The analysis targets four TG metrics: lag time, TTP, peak thrombin concentration, and ETP. For each disease group, 5,000 simulations were performed. The relative influence of each coagulation and inflammatory factor on these metrics for COVID-19 virtual patients was summarized in Figs. 5(A–D). Our results show that factor II concentration exerted the strongest positive effect on peak thrombin levels (Fig. 5(A)) and ETP (Fig. 5(D)), whereas ATIII showed the strongest negative association with both endpoints. Inflammatory mediators also displayed positive effects on peak and ETP, with IL-6 exhibiting a more pronounced influence than the other two inflammatory factors considered. Notably, although Factor VII is expected to positively regulate TG, PRCC analysis revealed a strong negative association with peak thrombin and ETP. This contradiction may arise from the structure of the Hockin model [29], in which excessive Factor VII leads to increased sequestration by TF, thereby limiting formation of the active VIIa–TF complex. The resulting reduction in effective VIIa–TF availability attenuates downstream coagulation cascade activation, ultimately diminishing thrombin peak and ETP. Consistent trends were observed for temporal TG metrics. Procoagulant and inflammatory factors predominantly exerted negative effects on TTP (Fig. 5(B)) and lag time (Fig. 5(C)), corresponding to accelerated TG, with Factor VIIa demonstrating the strongest time-shortening influence.

**Figure 5:**
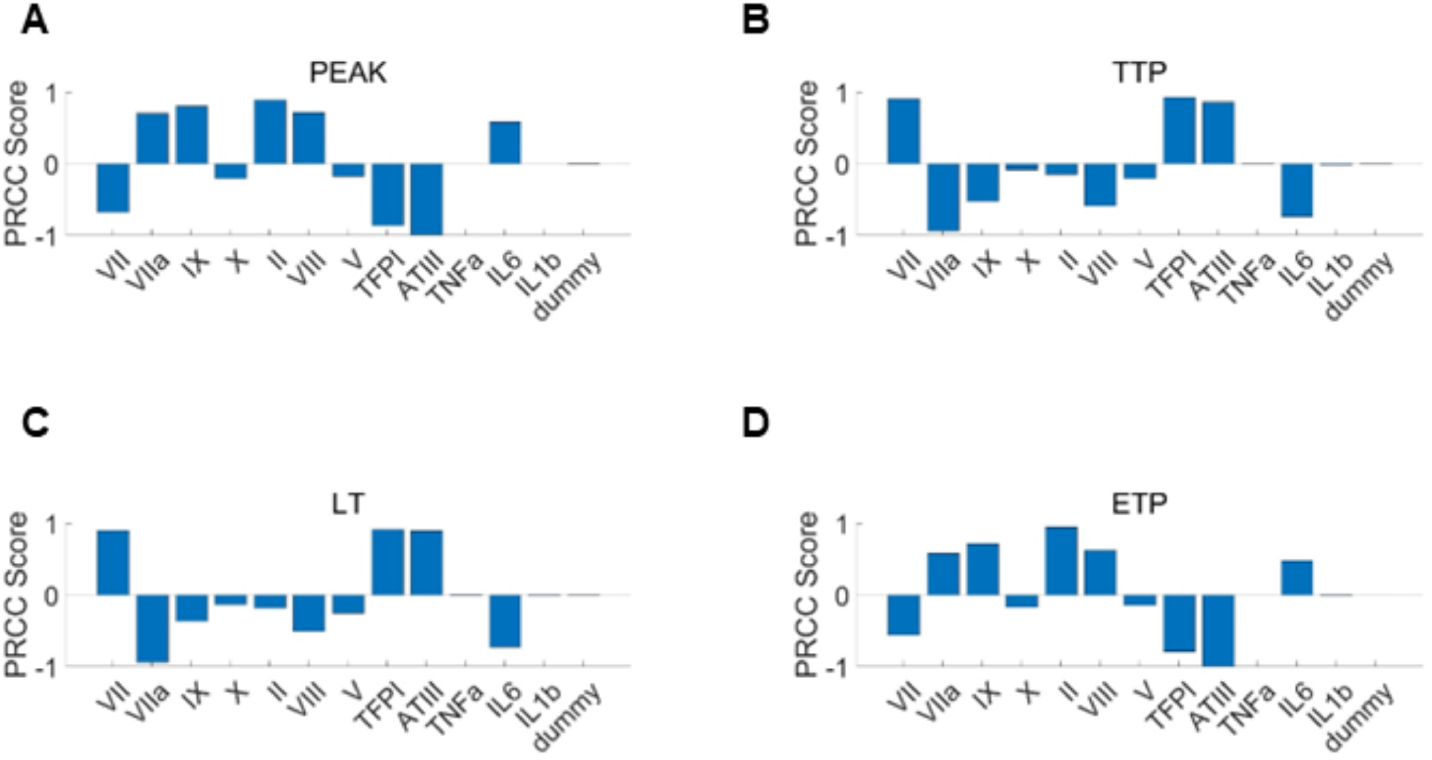
Sensitivity analysis of inflammatory and coagulation factors on the TG dynamics. Global sensitivity analysis using partial rank correlation coefficients quantified the influence of coagulation factors, anticoagulants, and inflammatory mediators on TG metrics, including Peak (A) time to peak (TTP, B), lag time (LT, C), and endogenous thrombin potential (ETP, D). The dummy variable acts as a control, representing a parameter with zero influence on the model output.

Based on these sensitivity analysis results, we narrowed the variation ranges of the three most influential coagulation factors when generating virtual patients for each disease such that the key TG metrics predicted align with the experimentally measured values reported in [53–56], while remaining within physiologically and clinically relevant bounds. Figs. 6(A-C) compare the simulated TG metrics against corresponding experimental measurements across the four examined patient groups, and the results show that simulated TG metrics generally fell within the experimental variability ranges for each disease group, demonstrating that the proposed virtual patient generation pipeline can quantitatively capture the observed TG dynamics across distinct disease states. These results will be used as baseline conditions for downstream analysis of the impact of pro-inflammatory cytokines.

**Figure 6:**
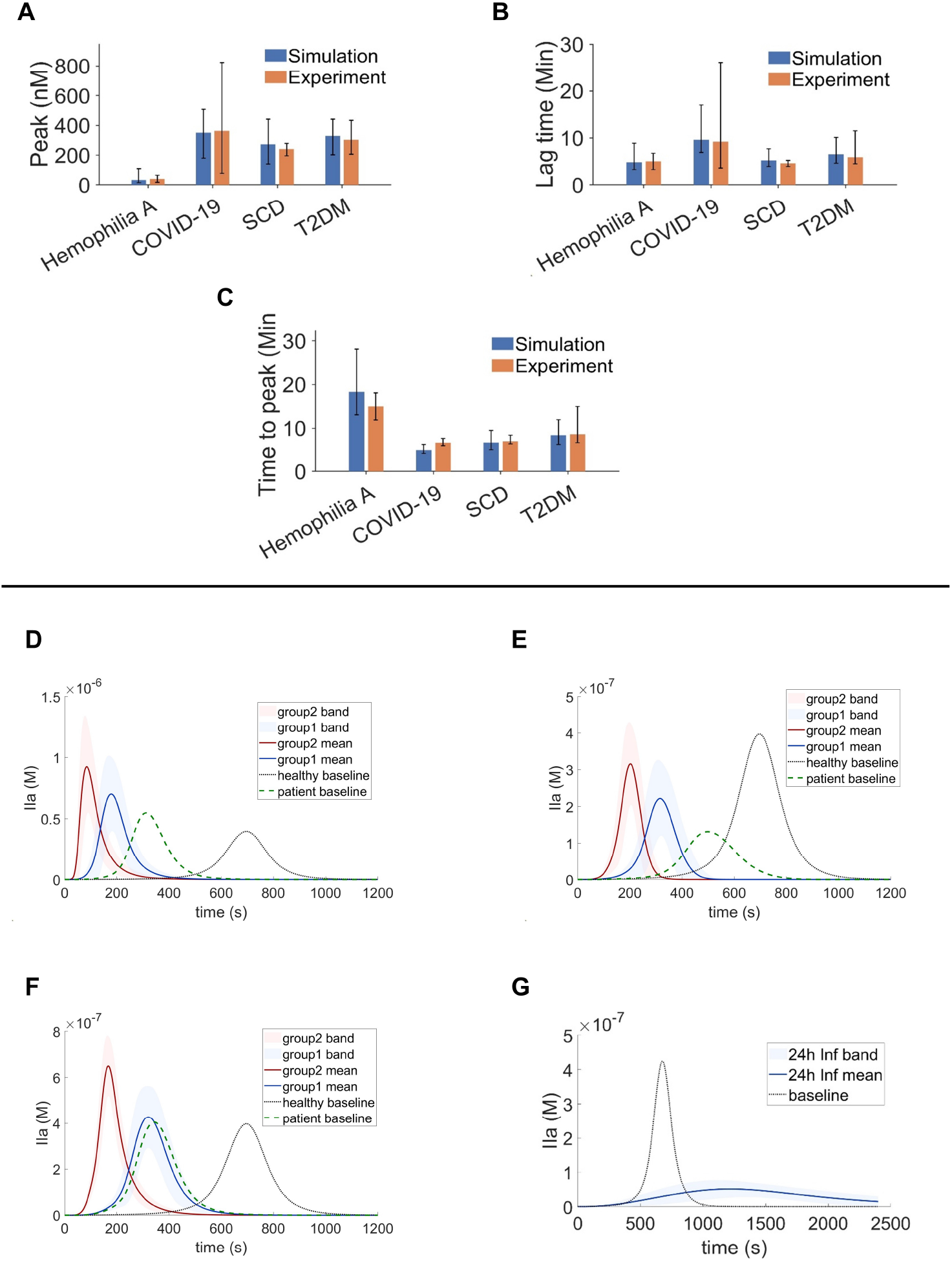
TG dynamics of virtual patients under four disease conditions. Comparison of simulated and experimental TG metrics, including peak (A), lag time (B), and time to peak (C), across hemophilia A, COVID-19, SCD, and T2DM. 2000 virtual patients are generated for each disease condition. TG profiles of virtual patients of COVID-19 (D), SCD (E), T2DM (F) under baseline conditions and inflammatory exposure (12 h and 24 h), shown as mean trajectories with variability bands. (G) TG profiles of virtual patients of Hemophilia A under baseline conditions. Virtual patient ensembles were constructed by sampling coagulation and inflammatory parameters within disease-specific physiological ranges.

Next, we examine the impact of pro-inflammatory cytokines on TG curves for virtual patient groups representing COVID-19 (Fig. 6(D)), SCD (Fig. 6(E)), and T2DM (Fig. 6(F)). Proinflammatory cytokine exposure was simulated at two durations, namely 12 hours and 24 hours, to assess how exposure length modulates coagulation response. For COVID-19 (Fig. 6(D)), both 12h and 24h inflammation exposures produce a dramatically elevated thrombin peak relative to both the uninflamed baseline and normal subjects, accompanied by shortened lag time and earlier time-to-peak. This pattern is consistent with the well-documented hypercoagulable state in COVID-19, driven by cytokine-mediated activation of coagulation pathways. Notably, the 24h exposure amplifies thrombin generation considerably beyond the 12h case, indicating a duration-dependent inflammatory response. In SCD (Fig. 6(E)), inflammation similarly shortens lag time and advances peak formation; however, the peak thrombin magnitude remains lower than that observed in normal subjects. This apparent paradox likely results from the chronically depleted plasma Factor X pool characteristic of SCD. Ongoing low-extent coagulation activity in SCD may consume Factor X at a rate that exceeds hepatic replenishment [83, 84], thereby constraining the maximum thrombin burst even under inflammatory stimulation.

For T2DM (Fig. 6(F)), the TG curves qualitatively mirror those of COVID-19, exhibiting short-ened lag time, earlier peak formation, and elevated TG peak relative to the uninflamed baseline and normal subjects, collectively indicating inflammation-driven enhancement of coagulation activity. Across all three conditions, the magnitude of these effects is mounted with exposure duration: prolonged inflammatory stimulation (24h) consistently produces earlier thrombin onset and higher peak IIa levels than shorter exposure (12h). In contrast to the above three diseases, the hemophilia A virtual patient population (Fig. 6(G)) exhibited a distinct response pattern. Under baseline conditions (without inflammatory signaling), TG remained markedly attenuated, consistent with a hypo-coagulable phenotype. This constrained thrombin amplification reflects a dominant effect imposed by coagulation factor VIII deficiency. Together, these results demonstrate that TG-calibrated virtual patient populations capture both inflammation-driven hypercoagulability mechanisms and disease-specific constraints on TG.

## 4 Discussion and Summary

In this study, we developed and calibrated a mechanistic inflammation-coagulation model and employed it in a virtual patient framework to investigate how inflammatory burden reshapes coagulation under hyperinflammatory conditions [9, 10, 85]. The model encodes cytokine-mediated upregulation of TF in monocytes and endothelial cells, concurrent suppression of endogenous anticoagulants, specifically ATIII and TFPI, and positive feedback from thrombin to inflammatory signaling [15, 86–89], which enables quantification of how the increasing inflammatory intensity and duration systematically alters TG at different disease conditions [15, 23, 90, 91]. Our results show that the virtual patient simulations reproduce trends qualitatively agree with clinical observations across different hypercoagulable diseases, including COVID-19, SCD and T2DM, demonstrating the feasibility of the model in simulating thrombo-inflammatory scenarios [23, 90–92]. Furthermore, our model shows that the interplay between inflammation and coagulation could promote a self-amplifying loop that leads to sustained thrombo-inflammatory conditions [85, 88]. This systems-level perspective provides a mechanistic explanation for why inflammatory states are frequently accompanied by heightened coagulation activity and thrombotic risk in severe inflammatory contexts [93, 94].

The inflammation intensity analysis in Figs. 4(A-C) highlights the nonlinear nature of inflammation–coagulation coupling. The increasing inflammatory burden leads to an exponential increase in TG, suggesting that coagulation activation is disproportionately sensitive to high cytokine levels. Importantly, virtual patient simulations demonstrate that both inflammatory intensity and duration critically impact the coagulation dynamics. Prolonged inflammatory stimulation resulted in earlier thrombin initiation and higher peak levels in COVID and T2DM patients compared with those with shorter exposure. These findings emphasize the importance of cumulative inflammatory burden rather than transient cytokine elevations, aligning with clinical observations linking sustained inflammation to progressive hypercoagulability [95, 96].

Disease-specific simulations revealed both shared and divergent TG dynamics under inflammatory conditions. In inflammatory diseases such as COVID-19 and T2DM, inflammatory signaling markedly amplified TG and accelerated coagulation initiation, consistent with clinical and experimental evidence of hypercoagulability in these settings [93, 97–99]. In contrast, the hemophilia A virtual population exhibited fundamentally attenuated TG dynamics, driven by factor VIII deficiency that constrains thrombin generation irrespective of inflammatory stimulus. This divergent behavior underscores the model’s capacity to capture a key mechanistic distinction between diseases, in which inflammation acts as the primary driver of coagulation dysregulation, and those in which an intrinsic factor deficiency suppresses the system-level coagulation activities.

Several limitations should be acknowledged. First, the model focuses on TG as a surrogate marker of coagulation activation without extending to the downstream consequences of thrombin activity, such as fibrin clot formation and propagation. Consequently, it cannot capture the full progression from coagulation initiation to thrombus formation and subsequent fibrinolysis. Second, the inflammatory module is built around a limited subset of cytokines, yet the *in vivo* inflammatory milieu is considerably more complex. Additional mediators, such as IL-12 and interferon-gamma (IFN-*γ*), have been reported to contribute to hypercoagulable conditions [100–102]. Due to this limitation, the model may underestimate or misrepresent inflammation–coagulation coupling in settings where these non-modeled mediators play a dominant role. Third, the TG data used for calibration of virtual patients were obtained from different laboratories employing distinct experimental protocols, which may introduce inter-laboratory variability that limits the feasibility of direct cross-disease comparison of thrombin generation dynamics.

In summary, this study introduces a mechanistic, TG-calibrated virtual patient framework that elucidates how inflammatory burden shapes coagulation dynamics. By incorporating interindividual variability, the proposed computational approach extends conventional TG assays to account for the interplay between inflammation and coagulation. This framework offers a potential tool for predicting disease progression, identifying disease-specific therapeutic targets, and supporting personalized management strategies in thrombo-inflammatory disorders.

## Acknowledgments

H.L. would like to thank Shan-Ho Tsai from the Georgia Advanced Computing Resource Center (GACRC) for providing technical support.

## References

[1] Monica Schenone, Barbara C Furie, and Bruce Furie. The blood coagulation cascade. Current opinion in hematology, 11(4):272–277, 2004.

[2] Robert B Francis. Clinical disorders of fibrinolysis: a critical review. Blut, 59(1):1–14, 1989.

[3] Bree Zimmerman and Leonard A Valentino. Hemophilia: in review. Pediatrics in review, 34(7):289–295, 2013.

[4] B Zoller, P Garcia de Frutos, Andreas Hillarp, and B Dahlback. Thrombophilia as a multigenic disease. Haematologica, 84(1):59–70, 1999.

[5] Konstantin Stark and Steffen Massberg. Interplay between inflammation and thrombosis in cardiovascular pathology. Nature Reviews Cardiology, 18(9):666–682, 2021.

[6] Charles T Esmon. Does inflammation contribute to thrombotic events? Haemostasis, 30(2):34–40, 2000.

[7] Paula A Klavina, Gemma Leon, Annie M Curtis, and Roger JS Preston. Dysregulated haemostasis in thrombo-inflammatory disease. Clinical Science, 136(24):1809–1829, 2022.

[8] Lei Zhu, He Dong, Lin Li, and Xiaojie Liu. The mechanisms of sepsis induced coagulation dysfunction and its treatment. Journal of inflammation research, pages 1479–1495, 2025.

[9] Charles T Esmon. The interactions between inflammation and coagulation. British Journal of Haematology, 131(4):417–430, 2005.

[10] Marcel Levi. Inflammation and coagulation. Inflammation: From Molecular and Cellular Mechanisms to the Clinic, pages 833–860, 2017.

[11] Marcel Levi, Tom van der Poll, and Harry R Buller. Bidirectional relation between inflammation and coagulation. Circulation, 109(22):2698–2704, 2004.

[12] Charles T Esmon, Kenji Fukudome, Tim Mather, Wolfram Bode, Lisa M Regan, Deborah J Stearns-Kurosawa, and Shinichiro Kurosawa. Inflammation, sepsis, and coagulation. Haematologica, 84(3):254–259, 1999.

[13] Marcel Levi and Tom van der Poll. Coagulation and sepsis. Thrombosis research, 149:38–44, 2017.

[14] Toshiaki Iba, Jerrold H Levy, Theodore E Warkentin, Jecko Thachil, Tom van der Poll, Marcel Levi, Scientific, Standardization Committee on DIC, the Scientific, Standardization Committee on Perioperative, Critical Care of the International Society on Thrombosis, and Haemostasis. Diagnosis and management of sepsis-induced coagulopathy and disseminated intravascular coagulation. Journal of Thrombosis and Haemostasis, 17(11):1989–1994, 2019.

[15] Guido Grignani and Anna Maiolo. Cytokines and hemostasis. Haematologica, 85(9):967–972, 2000.

[16] Michael Bode and Nigel Mackman. Regulation of tissue factor gene expression in monocytes and endothelial cells: thromboxane a2 as a new player. Vascular pharmacology, 62(2):57–62, 2014.

[17] C. Cerletti, C. Tamburrelli, B. Izzi, F. Gianfagna, and G. De Gaetano. Platelet-leukocyte interactions in thrombosis. Thrombosis Research, 129(3):263–266, 2012.

[18] Julie Rayes, Joshua H Bourne, Alexander Brill, and Steve P Watson. The dual role of platelet-innate immune cell interactions in thrombo-inflammation. Research and practice in thrombosis and haemostasis, 4(1):23–35, 2020.

[19] Shaun P Jackson, Roxane Darbousset, and Simone M Schoenwaelder. Thromboinflammation: challenges of therapeutically targeting coagulation and other host defense mechanisms. Blood, The Journal of the American Society of Hematology, 133(9):906–918, 2019.

[20] G. H. Frydman, A. Le, F. Ellett, J. Jorgensen, J. G. Fox, R. G. Tompkins, and D. Irimia. Technical advance: Changes in neutrophil migration patterns upon contact with platelets in a microfluidic assay. Journal of Leukocyte Biology, 101(3):797–806, 2017.

[21] Brittney Williams, Lin Zou, Jean-Francois Pittet, and Wei Chao. Sepsis-induced coagulopathy: a comprehensive narrative review of pathophysiology, clinical presentation, diagnosis, and management strategies. Anesthesia & Analgesia, 138(4):696–711, 2024.

[22] Andreas G Tsantes, Stavroula Parastatidou, Emmanuel A Tsantes, Elli Bonova, Konstantina A Tsante, Petros G Mantzios, Aristeidis G Vaiopoulos, Stavros Tsalas, Aikaterini Konstantinidi, Dimitra Houhoula, et al. Sepsis-induced coagulopathy: an update on pathophysiology, biomarkers, and current guidelines. Life, 13(2):350, 2023.

[23] Jean M Connors and Jerrold H Levy. Covid-19 and its implications for thrombosis and anticoagulation. Blood, The Journal of the American Society of Hematology, 135(23):2033– 2040, 2020.

[24] Douglas Chung, Suruchi Bakshi, and Piet H van der Graaf. A review of quantitative systems pharmacology models of the coagulation cascade: opportunities for improved usability. Pharmaceutics, 15(3):918, 2023.

[25] Andrew L Kuharsky and Aaron L Fogelson. Surface-mediated control of blood coagulation: the role of binding site densities and platelet deposition. Biophysical Journal, 80(3):1050– 1074, 2001.

[26] Aaron L Fogelson and Nessy Tania. Coagulation under flow: the influence of flow-mediated transport on the initiation and inhibition of coagulation. Pathophysiology of Haemostasis and Thrombosis, 34(2-3):91–108, 2005.

[27] Sharene D Bungay, Patricia A Gentry, and Rodney D Gentry. A mathematical model of lipid-mediated thrombin generation. Mathematical Medicine and Biology, 20(1):105–129, 2003.

[28] Manash S Chatterjee, Jeremy E Purvis, Lawrence F Brass, and Scott L Diamond. Pairwise agonist scanning predicts cellular signaling responses to combinatorial stimuli. Nature Biotechnology, 28(7):727–732, 2010.

[29] Matthew F Hockin, Kenneth C Jones, Stephen J Everse, and Kenneth G Mann. A model for the stoichiometric regulation of blood coagulation. Journal of Biological Chemistry, 277(21):18322–18333, 2002.

[30] Edward Beltrami and Jolyon Jesty. Mathematical analysis of activation thresholds in enzyme-catalyzed positive feedbacks: application to the feedbacks of blood coagulation. Proceedings of the National Academy of Sciences, 92(19):8744–8748, 1995.

[31] NM Dashkevich, MV Ovanesov, AN Balandina, SS Karamzin, PI Shestakov, NP Soshitova, AA Tokarev, MA Panteleev, and FI Ataullakhanov. Thrombin activity propagates in space during blood coagulation as an excitation wave. Biophysical Journal, 103(10):2233–2240, 2012.

[32] Ge Zhu, Susree Modepalli, Mohan Anand, and He Li. Computational modeling of hypercoagulability in COVID-19. Comput. Methods Biomech. Biomed. Engin., 26(3):338–349, February 2023.

[33] VI Zarnitsina, AV Pokhilko, and FI Ataullakhanov. A mathematical model for the spatiotemporal dynamics of intrinsic pathway of blood coagulation. i. the model description. Thrombosis research, 84(4):225–236, 1996.

[34] Mikhail A Panteleev, Mikhail V Ovanesov, Dmitrii A Kireev, Aleksei M Shibeko, Elena I Sinauridze, Natalya M Ananyeva, Andrey A Butylin, Evgueni L Saenko, and Fazoil I Ataullakhanov. Spatial propagation and localization of blood coagulation are regulated by intrinsic and protein c pathways, respectively. Biophysical journal, 90(5):1489–1500, 2006.

[35] Karin Leiderman and Aaron L Fogelson. Grow with the flow: a spatial–temporal model of platelet deposition and blood coagulation under flow. Mathematical Medicine and Biology: a Journal of the IMA, 28(1):47–84, 2011.

[36] Karin Leiderman and Aaron L Fogelson. The influence of hindered transport on the development of platelet thrombi under flow. Bulletin of Mathematical Biology, 75(8):1255–1283, 2013.

[37] Anass Bouchnita. Mathematical modelling of blood coagulation and thrombus formation under flow in normal and pathological conditions. PhD thesis, Université de Lyon; École Mohammadia d’ingénieurs (Rabat, Maroc), 2017.

[38] Anass Bouchnita, Kanishk Yadav, Jean-Pierre Llored, Alvaro Gurovich, and Vitaly Volpert. Thrombin generation thresholds for coagulation initiation under flow. Axioms, 12(9):873, 2023.

[39] Anass Bouchnita, Kirill Terekhov, Patrice Nony, Yuri Vassilevski, and Vitaly Volpert. A mathematical model to quantify the effects of platelet count, shear rate, and injury size on the initiation of blood coagulation under venous flow conditions. PloS one, 15(7):e0235392, 2020.

[40] Anass Bouchnita and Vitaly Volpert. A multiscale model of platelet-fibrin thrombus growth in the flow. Computers & Fluids, 184:10–20, 2019.

[41] A. Yazdani, H. Li, J. D. Humphrey, and G. E. Karniadakis. A general shear-dependent model for thrombus formation. PLoS Comput Biol, 13(1), 2017.

[42] Yazdani, H. Li, M. R. Bersi, P. D. Achille, J. Insley, J. D. Humphrey, and G. E. Karniadakis. Data-driven modeling of hemodynamics and its role on thrombus size and shape in aortic dissections. Sci Rep, 8, 2017.

[43] Yazdani, Y. Deng, H. Li, E. Javadi, Z. Li, S. Jamali, C. Lin, J. D. Humphrey, C. S. Mantzoros, and G.E. Karniadakis. Integrating blood cell mechanics, platelet adhesive dynamics and coagulation cascade for modelling thrombus formation in normal and diabetic blood. Journal of the Royal Society Interface, 18(175):20200834, 2021.

[44] He Li, Yixiang Deng, Zhen Li, Ander Dorken Gallastegi, Christos S Mantzoros, Galit H Frydman, and George E Karniadakis. Multiphysics and multiscale modeling of microthrombosis in covid-19. PLOS Computational Biology, 18(3):e1009892, 2022.

[45] Marco Witkowski, Ulf Landmesser, and Ursula Rauch. Tissue factor as a link between inflammation and coagulation. Trends in cardiovascular medicine, 26(4):297–303, 2016.

[46] Arthur J Chu. Tissue factor mediates inflammation. Archives of biochemistry and biophysics, 440(2):123–132, 2005.

[47] Kirk Johnson, Yoon Choi, Els DeGroot, Isa Samuels, Abla Creasey, and Lucien Aarden. Potential mechanisms for a proinflammatory vascular cytokine response to coagulation activation. The Journal of Immunology, 160(10):5130–5135, 1998.

[48] D Chen and A Dorling. Critical roles for thrombin in acute and chronic inflammation. Journal of Thrombosis and Haemostasis, 7:122–126, 2009.

[49] Paul M Ridker, Nader Rifai, Meir J Stampfer, and Charles H Hennekens. Plasma concentration of interleukin-6 and the risk of future myocardial infarction among apparently healthy men. Circulation, 101(15):1767–1772, 2000.

[50] Jan Klapproth, José Castell Thomas Geiger, Tilo Andus, and Peter C Heinrich. Fate and biological action of human recombinant interleukin 1β in the rat in vivo. European journal of immunology, 19(8):1485–1490, 1989.

[51] Rafael Simó, Anna Barbosa-Desongles, Albert Lecube, Cristina Hernandez, and David M Selva. Potential role of tumor necrosis factor-α in downregulating sex hormone–binding globulin. diabetes, 61(2):372–382, 2012.

[52] Johannes Schindelin, Curtis T Rueden, Mark C Hiner, and Kevin W Eliceiri. The imagej ecosystem: An open platform for biomedical image analysis. Molecular reproduction and development, 82(7-8):518–529, 2015.

[53] Omri Cohen, Nitsan Landau, Einat Avisahai, Tami Brutman-Barazani, Ivan Budnik, Tami Livnat, Keren Asraf, Ram Doolman, Sarina Levy-Mendelovich, Orly Efros, et al. Association between thrombin generation and clinical characteristics in covid-19 patients. Acta Haematologica, 146(2):151–160, 2023.

[54] Armando Tripodi, Adriana Branchi, Veena Chantarangkul, Marigrazia Clerici, Giuliana Merati, Andrea Artoni, and Pier Mannuccio Mannucci. Hypercoagulability in patients with type 2 diabetes mellitus detected by a thrombin generation assay. Journal of thrombosis and thrombolysis, 31(2):165–172, 2011.

[55] Chirag Amin, Soheir Adam, Micah J Mooberry, Abdullah Kutlar, Ferdane Kutlar, Denise Esserman, Julia E Brittain, Kenneth I Ataga, Jen-Yea Chang, Alisa S Wolberg, et al. Coagulation activation in sickle cell trait: an exploratory study. British journal of haematology, 171(4):638–646, 2015.

[56] Romy MW de Laat-Kremers, Marisa Ninivaggi, Iris van Moort, Moniek de Maat, and Bas de Laat. Tailoring the effect of antithrombin-targeting therapy in haemophilia a using in silico thrombin generation. Scientific Reports, 11(1):15572, 2021.

[57] D. M. Del Valle, S. Kim-Schulze, H. Huang, N. D. Beckmann, S. Nirenberg, B. Wang, Y. Lavin, T. H. Swartz, D. Madduri, A. Stock, et al. An inflammatory cytokine signature predicts COVID-19 severity and survival. Nature Medicine, 26(10):1636–1643, 2020.

[58] S Mirza, Monir Hossain, Christine Mathews, Perla Martinez, Paula Pino, Jennifer L Gay, Anne Rentfro, Joseph B McCormick, and Susan P Fisher-Hoch. Type 2-diabetes is associated with elevated levels of tnf-alpha, il-6 and adiponectin and low levels of leptin in a population of mexican americans: a cross-sectional study. Cytokine, 57(1):136–142, 2012.

[59] Stephen C Taylor, Samuel J Shacks, Ralph A Mitchell, and Aaron Banks. Serum interleukin-6 levels in the steady state of sickle cell disease. Journal of interferon & cytokine research, 15(12):1061–1064, 1995.

[60] Inés Malavé, Yolanda Perdomo, Edgar Escalona, Ernesto Rodriguez, Miren Anchustegui, Hector Malavé, and Tulio Arends. Levels of tumor necrosis factor α/cachectin (tnfα) in sera from patients with sickle cell disease. Acta haematologica, 90(4):172–176, 1993.

[61] Liliane K Siransy, Romuald S Dasse, Honore Adou, Patricia Kouacou, Sidonie Kouame-nan, Yassongui Sekongo, Richard Yeboah, Charlene Memel, Aniella Assi-Sahoin, Salimata Y Moussa, et al. Are il-1 family cytokines important in management of sickle cell disease in sub-saharan africa patients? Frontiers in Immunology, 14:954054, 2023.

[62] Bingwen Eugene Fan, Jensen Ng, Stephrene Seok Wei Chan, Dheepa Christopher, Allison Ching Yee Tso, Li Min Ling, Barnaby Edward Young, Lester Jun Long Wong, Christina Lai Lin Sum, Hwee Tat Tan, et al. Covid-19 associated coagulopathy in critically ill patients: a hypercoagulable state demonstrated by parameters of haemostasis and clot waveform analysis. Journal of thrombosis and thrombolysis, 51(3):663–674, 2021.

[63] Sanjeev Palta, Richa Saroa, and Anshu Palta. Overview of the coagulation system. Indian Journal of Anaesthesia, 58(5):515, 2014.

[64] B Nsiri, N Gritli, F Bayoudh, T Messaoud, S Fattoum, and S Machghoul. Abnormalities of coagulation and fibrinolysis in homozygous sickle cell disease. Hematology and cell therapy, 38(3):279–284, 1996.

[65] Rehab S El-Hagracy, Gihan M Kamal, Inas M Sabry, Abeer A Saad, Nahla F Abou El Ezz, and Hesham AR Nasr. Tissue factor, tissue factor pathway inhibitor and factor vii activity in cardiovascular complicated type 2 diabetes mellitus. Journal of Diabetology, 1(1):4, 2010.

[66] Pratima Chowdary. Inhibition of tissue factor pathway inhibitor (tfpi) as a treatment for haemophilia: Rationale with focus on concizumab: P. chowdary. Drugs, 78(9):881–890, 2018.

[67] Diapharma. Antithrombin (atiii), 2024.

[68] Alok Srivastava, Elena Santagostino, Alison Dougall, Steve Kitchen, Megan Sutherland, Steven W Pipe, Manuel Carcao, Johnny Mahlangu, Margaret V Ragni, Jerzy Windyga, et al. Wfh guidelines for the management of hemophilia. Haemophilia, 26:1–158, 2020.

[69] Saulius Butenas, Cornelis van’t Veer, and Kenneth G Mann. “normal” thrombin generation: Presented in part at the xvith congress of the international society on thrombosis and haemostasis, june 6-12, 1997, Florence, Italy (abstr ps-1653), at the 15th international congress on thrombosis, october 16-21, 1998, antalya, turkey (abstr 234), and at the 40^th^ annual meeting of the american society of hematology, december 4-8, 1998, miami beach, fl (abstr 151). Blood, The Journal of the American Society of Hematology, 94(7):2169–2178, 1999.

[70] Giovanni Cimmino, Stefano Conte, Mariarosaria Morello, Grazia Pellegrino, Laura Marra, Andrea Morello, Giuseppe Nicoletti, Gennaro De Rosa, Paolo Golino, and Plinio Cirillo. Vitamin d inhibits il-6 pro-atherothrombotic effects in human endothelial cells: a potential mechanism for protection against covid-19 infection? Journal of Cardiovascular Development and Disease, 9(1):27, 2022.

[71] Franz-Josef Neumann, Ilka Ott, Nikolaus Marx, Thomas Luther, Silke Kenngott, Meinrad Gawaz, Mathias Kotzsch, and Albert Schomig. Effect of human recombinant interleukin-6 and interleukin-8 on monocyte procoagulant activity. Arteriosclerosis, thrombosis, and vascular biology, 17(12):3399–3405, 1997.

[72] Markus Puhlmann, David M Weinreich, Jeffrey M Farma, Nancy M Carroll, Ewa M Turner, and H Richard Alexander Jr. Interleukin-1β induced vascular permeability is dependent on induction of endothelial tissue factor (tf) activity. Journal of translational medicine, 3(1):37, 2005.

[73] Antje Scholl, Igor Ivanov, and Burkhard Hinz. Inhibition of interleukin-1β-induced endothelial tissue factor expression by the synthetic cannabinoid win 55,212-2. Oncotarget, 7(38):61438, 2016.

[74] Anna K Brandtner, Georg F Lehner, Andreas Pircher, Clemens Feistritzer, and Michael Joannidis. Differential procoagulatory response of microvascular, arterial and venous endothelial cells upon inflammation in vitro. Thrombosis research, 205:70–80, 2021.

[75] Jihwa Chung, Takatoshi Koyama, Mai Ohsawa, Aya Shibamiya, Asuka Hoshi, and Shinsaku Hirosawa. 1, 25 (oh) 2d3 blocks tnf-induced monocytic tissue factor expression by inhibition of transcription factors ap-1 and nf-κb. Laboratory investigation, 87(6):540–547, 2007.

[76] RWLM Niessen, RJ Lamping, PM Jansen, MH Prins, M Peters, FB Taylor Jr, JJM De Vijlder, JW Ten Cate, CE Hack, and A Sturk. Antithrombin acts as a negative acute phase protein as established with studies on hepg2 cells and in baboons. Thrombosis and haemostasis, 78(09):1088–1092, 1997.

[77] Hong Jin, Wen-Bing Qiu, Yi-Fang Mei, Dong-Ming Wang, Yu-Guang Li, and Xue-Rui Tan. Testosterone alleviates tumor necrosis factor-alpha-mediated tissue factor pathway inhibitor downregulation via suppression of nuclear factor-kappa b in endothelial cells. Asian journal of andrology, 11(2):266, 2009.

[78] Tomotake Tokunou, Toshihiro Ichiki, Kotaro Takeda, Yuko Funakoshi, Naoko Iino, Hiroaki Shimokawa, Kensuke Egashira, and Akira Takeshita. Thrombin induces interleukin-6 expression through the camp response element in vascular smooth muscle cells. Arteriosclerosis, thrombosis, and vascular biology, 21(11):1759–1763, 2001.

[79] Grzegorz Wilhelm, Paulina Mertowska, Sebastian Mertowski, Anna Przysucha, Jerzy Strużyna, Ewelina Grywalska, and Kamil Torres. The crossroads of the coagulation system and the immune system: interactions and connections. International journal of molecular sciences, 24(16):12563, 2023.

[80] Rasoul Ebrahimi, Fatemeh Nasri, and Tahereh Kalantari. Coagulation and inflammation in covid-19: reciprocal relationship between inflammatory and coagulation markers. Annals of Hematology, 103(6):1819–1831, 2024.

[81] Rachelle P Davis, Sarah Miller-Dorey, and Craig N Jenne. Platelets and coagulation in infection. Clinical & Translational Immunology, 5(7):e89, 2016.

[82] Simeone Marino, Ian B Hogue, Christian J Ray, and Denise E Kirschner. A methodology for performing global uncertainty and sensitivity analysis in systems biology. Journal of theoretical biology, 254(1):178–196, 2008.

[83] Denis Noubouossie, Nigel S Key, and Kenneth I Ataga. Coagulation abnormalities of sickle cell disease: Relationship with clinical outcomes and the effect of disease modifying therapies. Blood reviews, 30(4):245–256, 2016.

[84] Erica Sparkenbaugh and Rafal Pawlinski. Interplay between coagulation and vascular inflammation in sickle cell disease. British Journal of Haematology, 162(1):3–14, 2013.

[85] Bernd Engelmann and Steffen Massberg. Thrombosis as an intravascular effector of innate immunity. Nature Reviews Immunology, 13(1):34–45, 2013.

[86] JM Herbert, P Savi, MC Laplace, and A Lale. Il-4 inhibits lps-, il-1β-and tnfα-induced expression of tissue factor in endothelial cells and monocytes. FEBS letters, 310(1):31–33, 1992.

[87] Alireza R Rezaie. Protease-activated receptor signalling by coagulation proteases in endothelial cells. Thrombosis and haemostasis, 112(11):876–882, 2014.

[88] Shaun R Coughlin. Thrombin signalling and protease-activated receptors. Nature, 407(6801):258–264, 2000.

[89] Melissa E Schechter, Bruno B Andrade, Tianyu He, George Haret Richter, Kevin W Tosh, Benjamin B Policicchio, Amrit Singh, Kevin D Raehtz, Virginia Sheikh, Dongying Ma, et al. Inflammatory monocytes expressing tissue factor drive siv and hiv coagulopathy. Science translational medicine, 9(405):eaam5441, 2017.

[90] Denis F Noubouossie, Phu Quoc Lê, Francis Corazza, France Debaugnies, Laurence Rozen, Alina Ferster, and Anne Demulder. Thrombin generation reveals high procoagulant potential in the plasma of sickle cell disease children. American journal of hematology, 87(2):145–149, 2012.

[91] Marieke JA Verhagen, Waander L van Heerde, Johanna G van der Bom, Erik AM Beckers, Nicole MA Blijlevens, Michiel Coppens, Samantha C Gouw, Joop H Jansen, Frank WG Leebeek, Lize FD van Vulpen, et al. In patients with hemophilia, a decreased thrombin generation profile is associated with a severe bleeding phenotype. Research and practice in thrombosis and haemostasis, 7(2):100062, 2023.

[92] Hui Yin Lim, Brandon Lui, Mark Tacey, Anna Kwok, Suresh Varadarajan, Geoffrey Donnan, Harshal Nandurkar, and Prahlad Ho. Global coagulation assays in patients with diabetes mellitus. Research and Practice in Thrombosis and Haemostasis, 5(7):e12611, 2021.

[93] Edward M Conway, Nigel Mackman, Ronald Q Warren, Alisa S Wolberg, Laurent O Mosnier, Robert A Campbell, Lisa E Gralinski, Matthew T Rondina, Frank L van de Veerdonk, Karin M Hoffmeister, et al. Understanding covid-19-associated coagulopathy. Nature Reviews Immunology, 22(10):639–649, 2022.

[94] Behnood Bikdeli, Mahesh V Madhavan, David Jimenez, Taylor Chuich, Isaac Dreyfus, Elissa Driggin, Caroline Der Nigoghossian, Walter Ageno, Mohammad Madjid, Yutao Guo, et al. Covid-19 and thrombotic or thromboembolic disease: implications for prevention, antithrombotic therapy, and follow-up: Jacc state-of-the-art review. Journal of the American college of cardiology, 75(23):2950–2973, 2020.

[95] Shehan N Randeria, Greig JA Thomson, Theo A Nell, Timothy Roberts, and Etheresia Pretorius. Inflammatory cytokines in type 2 diabetes mellitus as facilitators of hypercoagulation and abnormal clot formation. Cardiovascular diabetology, 18(1):72, 2019.

[96] Andrea Boccatonda, Elena Campello, Chiara Simion, and Paolo Simioni. Long-term hypercoagulability, endotheliopathy and inflammation following acute sars-cov-2 infection. Expert review of hematology, 16(12):1035–1048, 2023.

[97] Nicola Conran and Erich V De Paula. Thromboinflammatory mechanisms in sickle cell disease–challenging the hemostatic balance. Haematologica, 105(10):2380, 2020.

[98] Giuseppe Patti, Ilaria Cavallari, Felicita Andreotti, Paolo Calabro, Plinio Cirillo, Gentian Denas, Mattia Galli, Enrica Golia, Ernesto Maddaloni, Rossella Marcucci, et al. Prevention of atherothrombotic events in patients with diabetes mellitus: from antithrombotic therapies to new-generation glucose-lowering drugs. Nature Reviews Cardiology, 16(2):113–130, 2019.

[99] Rowena Brook, Mani Suleiman, Brandon Lui, Ramzi A Ajjan, Harshal Nandurkar, Suresh Varadarajan, Prahlad Ho, and Hui Yin Lim. Global coagulation assays for patients with diabetes and their utility in the prediction of atherothrombotic events. Blood Vessels, Thrombosis & Hemostasis, 2(3):100079, 2025.

[100] Junko Kato, Tomohiro Okamoto, Hiroyuki Motoyama, Ryosuke Uchiyama, Daniel Kirchhofer, Nico Van Rooijen, Hirayuki Enomoto, Shuhei Nishiguchi, Norifumi Kawada, Jiro Fujimoto, et al. Interferon-gamma–mediated tissue factor expression contributes to t-cell-mediated hepatitis through induction of hypercoagulation in mice. Hepatology, 57(1):362–372, 2013.

[101] J.E.A. Portielje, W. H.J. Kruit, A. J.M. Eerenberg, M. Schuler, A. Sparreboom, C. H.J. Lamers, R.L.H. Bolhuis, G. Stoter, C. Huber, and C. E. Hack. Interleukin 12 induces activation of fibrinolysis and coagulation in humans. British Journal of Haematology, 112(2):499– 505, 2001.

[102] Martin J Page, Janette Bester, and Etheresia Pretorius. Interleukin-12 and its procoagulant effect on erythrocytes, platelets and fibrin (ogen): the lesser known side of inflammation. British journal of haematology, 180(1):110–117, 2018.

